# Non-invasive disruption of the blood-brain barrier in the marmoset monkey

**DOI:** 10.1101/2022.11.08.515696

**Authors:** Diego Szuzupak, Sang-Ho Choi, Aydin Alikaya, Yongshan Mou, Afonso C. Silva, David J. Schaeffer

## Abstract

The common marmoset monkey (*Callithrix jacchus*) is a species of rising prominence in the neurosciences due to their small size, ease of handling, fast breeding, and their shared functional and structural brain characteristics with Old World primates. With increasing attention on modeling human brain diseases in marmosets, understanding how to deliver therapeutic or neurotropic agents to the marmoset brain non-invasively is of great preclinical importance. In other species, including humans, transcranial focused ultrasound (tFUS) aided by intravenously injected microbubbles has proven to be a transient, reliable, and safe method for disrupting the blood-brain barrier (BBB), allowing for the focal passage of therapeutic agents that do not otherwise readily traverse the tight endothelial junctions of the BBB. The critical gap that we address here is to document parameters to disrupt the BBB reliably and safely in marmosets using tFUS. By integrating our marmoset brain atlases and the use of a marmoset-specific stereotactic targeting system, we conducted a series of systematic transcranial sonication experiments in nine marmosets. We demonstrate the effects of center frequency, acoustic pressure, burst period and duration, establish a minimum microbubble dose, estimate microbubble clearance time, and estimate the duration that the BBB remained open to passage. Successful BBB disruption was reported *in vivo* with MRI-based contrast agents, as well as Evans blue staining assessed *ex vivo*. Histology (Hematoxylin and Eosin staining) and immunohistochemistry indicated that the BBB can be safely and reliably opened with the parameters derived from these experiments.

## Introduction

The focus of this manuscript is to establish parameters to safely disrupt the blood-brain barrier (BBB) in the common marmoset monkey (*Callithrix jacchus*), a species of rising prominence in the neurosciences. The BBB regulates the permeability of molecules to the brain parenchyma, consisting of capillary endothelium that prevents molecules with a weight of ~400 Da from entering^1^. In other preclinical modelling species (e.g., rats, mice, macaques, rabbits, pigs), transcranial focused ultrasound (tFUS) has become a reliable means to circumvent invasive intracerebral injections and allow for agent delivery by transiently disrupting the BBB^2–9^. The value of applying tFUS to non-invasively and locally disrupt the BBB in the marmoset model is potentially tremendous – for example, by taking advantage of the marmoset’s short interbirth interval and relatively short lifespan, tFUS can be used as a longitudinal and non-invasive method of neuromodulation^10^, neuronal tracing^11,12^, or even focal drug delivery^13^ for a disease model across the lifespan. With a lissencephalic cortex and cortical architecture that is more similar to humans than to rodents^14^, marmosets are ideal for tFUS, allowing for simplified targeting across the cortical ribbon when compared to the highly folded brains of other primate species.

The critical gap addressed here is to document the ability to reliably and non-invasively open the BBB in the marmoset using tFUS aided by microbubble cavitation. Microbubbles are microscopic (~1-10 μm) gas-filled micelles that can be systemically injected just before ultrasonic stimulation^15^. At lower acoustic pressures, microbubbles oscillate (stable cavitation) and when exposed to sufficient pressure can collapse (inertial cavitation) and release a powerful liquid jet through the endothelium that can potentiate agent delivery^15–17^. With myriad available neuromodulatory or therapeutic agents having been shown to cross the BBB as a result of ultrasound-mediated microbubble cavitation^13,18–20^, understanding the parameters to open the BBB in the marmoset will inevitably lead to numerous neuroscientific applications. We expect this technique will have broad application in translational marmoset research, especially for neurodevelopmental applications, providing a means to track the neuropathological emergence of circuit dysfunction non-invasively from a young age.

Through the combination of a marmoset-specific atlas^21–24^ and a now commercially available marmoset-specific focused ultrasound apparatus (Figure 1; RK-50 Marmoset, FUS Instruments Incorporated, Toronto, ON, Canada), we demonstrate the requisite parameters to reliably stereotactically target and open the BBB to a small parenchymal volume (~10 – 20 mm^3^) with a single element 1.46 MHz transducer. We demonstrate the effects of center frequency on BBB disruption size, the minimum acoustic pressure to disrupt the BBB, and the deleterious effects of too much acoustic pressure. We also demonstrate the minimum microbubble dosage for BBB disruption at safe acoustic pressure, estimate microbubble clearance time, and the effect of skull angle on BBB disruption. Through reporting by *in vivo* gadolinium- and manganese chloride-enhanced MRI and *ex vivo* Evans Blue staining as well as histological and immunohistochemical reporting, we provide a detailed account of the requisite parameters for safely, reproducibly, and focally disrupting the BBB in the common marmoset.

**Figure 1.**
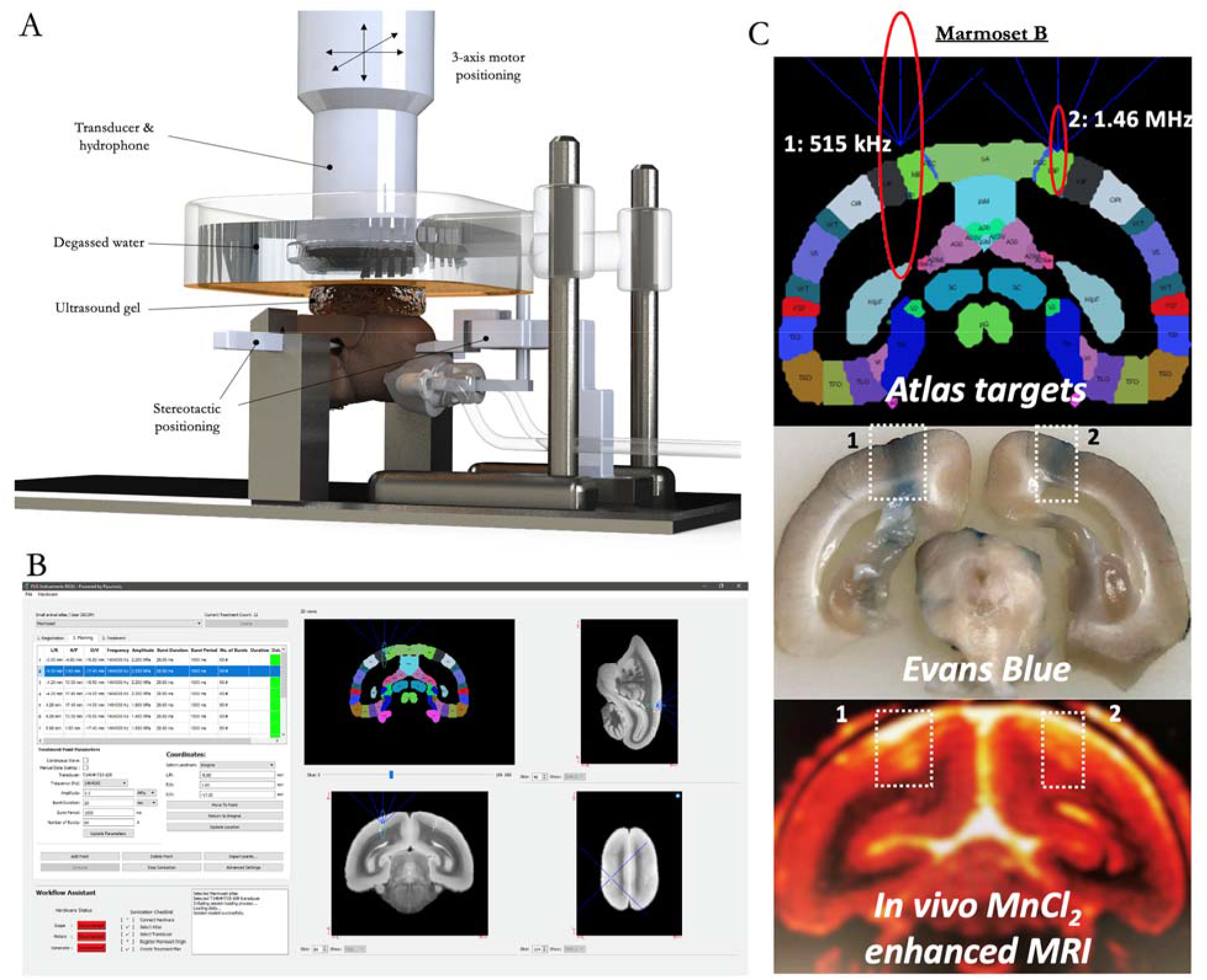
Focused ultrasound positioning apparatus, customized for use in marmoset monkeys. (A) shows computer-aided design 3-dimensional rendering of the stereotactic positioning of the marmoset with reference to the transducer and hydrophone. (B) shows the software MORPHEUS (FUS Instruments, Toronto, Ontario, Canada) with our marmoset atlases integrated along with cytoarchitectonic boundaries. (C) demonstrates the relative size of BBB disruption using 515 kHz and 1.46 MHz transducers, as reported by Evans blue staining and manganese enhanced MRI.

## Methods

### Animals

Nine adult marmosets (*Callithrix jacchus*) contributed data to this study (Table 1). Animals were anesthetized (induced and maintained) with 2 % isoflurane delivered via mask for both the tFUS and *in vivo* MRI procedures. During the procedures, heart rate, blood oxygenation, respiration, and rectal temperature were monitored. The head was shaved with clippers, then any remaining hair was removed with depilatory cream. A 26-gauge catheter (1/2- or 3/4-inch length) was placed in the lateral tail or saphenous vein for microbubble and contrast agent delivery. Body temperature was maintained with infrared or heated water blankets. Experimental procedures were approved by the University of Pittsburgh Institutional Animal Care and Use Committee.

**Table 1.** Summary of animal age, weight, and sex.

|  | **Age (months)** | **Weight (grams)** | **Sex** |
| --- | --- | --- | --- |
| Marmoset B | 46 | 205 | F |
| Marmoset SP | 32 | 195 | F |
| Marmoset SG | 24 | 135 | F |
| Marmoset SK | 20 | 170 | F |
| Marmoset NE | 29 | 175 | M |
| Marmoset T | 19 | 145 | F |
| Marmoset E | 18 | 165 | M |
| Marmoset G | 40 | 215 | F |
| Marmoset M | 97 | 235 | M |

### Focused ultrasound apparatus

Sonications were performed with the RK-50 Marmoset (FUS Instruments Incorporated, Toronto, ON, Canada), which was developed for use in marmosets in collaboration with FUS Instruments to implement marmoset-specific hardware, including a marmoset stereotaxic device (Model SR-AC; Narishige International Incorporated, Amityville, New York, USA) and to include an MRI-based marmoset atlas for stereotactic targeting^21,22^ (Figure 1). The “turn-key” device features an automatic 3-axis positioning system that is configured with reference to stereotactic position on the treatment planning workstation. Two 35 mm spherically focused and calibrated transducers were used in this study, a 515 kHz and a 1.46 MHz transducer (FUS Instruments Incorporated, Toronto, ON, Canada), driven by a 15 W amplifier. A digital oscilloscope was used to monitor cavitation emissions from the hydrophone, which was mounted concentrically with the transducers. As illustrated in Figure 1A, the transducer was sealed with a 3D printed cover with an o-ring seal and a polyimide film face that held degassed water (via portable water degasser; FUS-DS-50, FUS Instruments Incorporated, Toronto, ON, Canada). The sensor and cover were immersed in a tank holding ~300 ml of degassed water. The polyimide base of the tank was coupled with the head via ultrasound gel.

### Stereotactic atlas-based targeting

Sonications were applied transcranially (with skin intact, only hair removed) based on stereotactic position, with x = 0, y = 0, z = 0 mm corresponding to midway between the center of the ear bars, in plane with the bottom of the orbit bars. The marmoset-specific stereotax was rigidly mounted the RK-50 Marmoset baseplate with thumbscrews. Sonication sites were then specified based on the desired location in stereotactic space (Figure 1B for example). For co-localization with functional and structural atlas landmarks in the marmoset brain, the tFUS positioning software (MORPHEUS framework, FUS Instruments Incorporated, Toronto, ON, Canada) was integrated with the marmosetbrainmapping.org^25^ and marmosetbrainconnectome.org^22^ atlases. Together with the cytoarchitectonic boundaries derived from the Paxinos and RIKEN marmoset brain atlases^24,26^, the Morpheus software allowed for accurate positioning of the transducer with reference to the adult marmoset brain.

### Sonications

#### Microbubbles

Immediately prior to the sonication (<1 minute) microbubbles (Definity, Lantheus Medical Imaging, Billerica, MA, USA), were administered via lateral tail or saphenous vein catheter to aide in BBB disruption. Microbubble solutions were injected directly into the catheter hub; a 26-gauge catheter was chosen to reduce the probability of premature microbubble destruction. The concertation (20 μL/kg – 400 μL/kg) of microbubbles varied across experiments (detailed in turn), but all injections were prepared in a stock solution (100 μL microbubbles / 860 μL sterile saline) in a 1 mL syringe (weight (kg) x microbubble concertation (mL/kg) x 9.6 = injection volume (mL)) for all experiments. The solution was injected as a bolus and flushed with 200 μL of sterile saline to ensure that the microbubbles cleared the volume of the catheter hub.

#### Evans blue & MRI contrast agent injections

Evans blue stain and one of two MRI contrast agents (manganese chloride or gadolinium) were used to verify BBB disruption. All agents were injected immediately after the last sonication as a bolus (albeit the injection timing varied for the BBB disruption duration and clearance experiments, detailed below). Evans blue (E2129; Sigma-Aldrich Co., MO) was injected intravenously as a bolus in all animals at a dose of 2 μL/g in 2 % solution, prepared in sterile water. Manganese chloride (M5005; Sigma-Aldrich Co., MO) at a concentration of 100 mM was buffered in a 400 mM solution of Bicine (B3876; Sigma-Aldrich Co., MO), corrected to a pH of 7.4 with a 1 M sodium hydroxide solution^27^ (655104; Sigma-Aldrich Co., MO) and filtered with a sterile syringe filter at 2 μm. 200 μL of MnCl2 solution was injected in Marmoset B only, thereafter a gadolinium base contrast agent (GBCA) was used for *in vivo* MRI contrast. Gadolinium (Gadavist™, gadobutrol; Bayer Healthcare Pharmaceuticals, Leverkusen, Germany) was prepared in 200 μL of sterile saline and injected at a dose of 100 μL/kg in all marmosets except Marmosets B (MnCl2 used), G (200 μL/kg), and SP (600 μL/kg).

#### BBB disruption as a function of center frequency

Marmoset B received two sonications in parietal cortices to determine the relative difference in BBB disruption size as a function of transducer center frequency (Figure 1C). Left parietal cortex was sonicated with a 515 kHz transducer (1 MPa, 30 ms burst duration, 1,000 ms burst period, 90 bursts, 400 μL/kg microbubble dose). Right parietal cortex was sonicated with a 1.46 MHz transducer (3.2 MPa, 30 ms burst duration, 1,000 ms burst period, 90 bursts, 400 μL/kg microbubble dose). BBB disruption was reported by *in vivo* manganese-enhanced MRI and Evans blue staining *ex vivo*.

#### BBB disruption as a function of acoustic pressure

For this experiment, marmosets SG, NE, and T each received five or eight cortical sonications (area 8a, 4ab, MIP, and V2) with a 1.46 MHz transducer. Figure 2 shows the sonication locations and accompanying acoustic pressures. For Marmoset SG, the acoustic pressure was modulated with the following parameters remaining constant: burst duration = 30 ms, burst period = 1,000 ms, number of bursts = 90, microbubble dose = 400 μL/kg. Based on the information gained from Marmoset SG, Marmosets NE and T were sonicated at lower acoustic pressures with the following parameters remaining constant: burst duration = 20 ms, burst period = 1,000 ms, number of bursts = 60, microbubble dose = 200 μL/kg. The purpose of this experiment was threefold: to (1) determine the minimum acoustic pressure to open the BBB in a marmoset at 1.46 MHz, (2) determine the size/shape of BBB disruption as a function of acoustic pressure, and (3) determine the pressure at which damage starts to occur. For all three animals, BBB disruption was reported by *in vivo* GBCA-enhanced MRI and Evans blue staining *ex vivo*. Tissue damage was assessed using H & E staining (detailed below).

**Figure 2.**
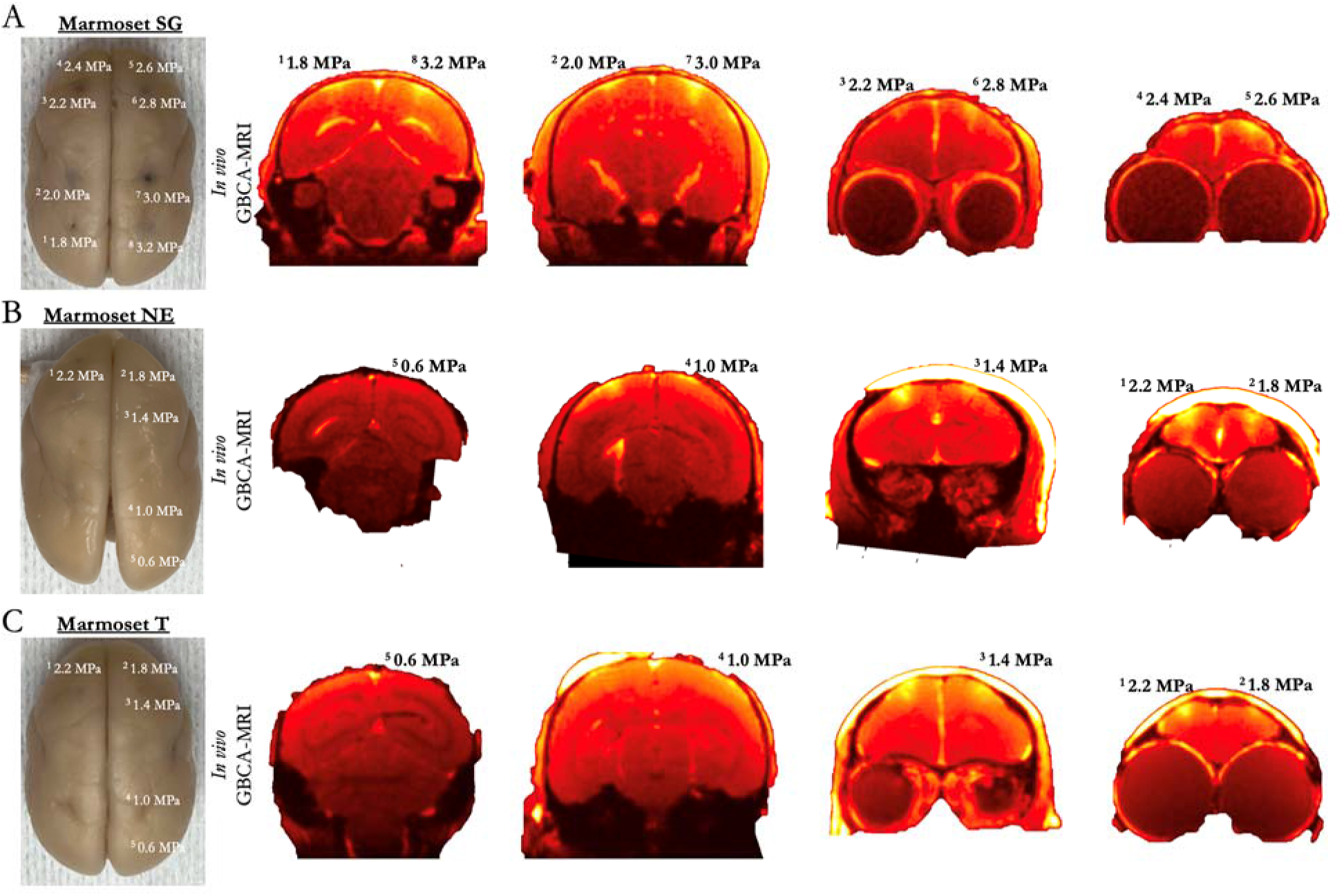
Blood-brain barrier disruption as a function of acoustic pressure, minimum pressure. (A) shows the resultant disruption (extravasation of Evans blue, left photo; and GBCA using *in vivo* MRI, right) of the BBB at 8 sonications sites across the cortex of marmoset SG. Panels B and C shows extravasation in the same manner as panel A, but in marmosets NE and T, which were sonicated at lower acoustic pressures.

#### Minimum microbubble dosage to open the BBB

Marmosets NE and T were used to determine the minimum microbubble dosage necessary to open the BBB. Each animal was sonicated four times (area 8a, 4ab, MIP, and V2), with differing microbubble dosages from 0 to 200 μL/kg (Figure 3 shows dosages at specific locations), with the following parameters remaining constant: acoustic pressure = 2.2 MPa, burst duration = 20 ms, burst period = 1,000 ms, number of bursts = 60. Ten minutes elapsed between sonications and microbubble injections to reduce the confounding effects of circulating microbubbles. Note that Marmosets NE and T participated in both the acoustic pressure (above) and microbubble dosing experiment (albeit separate sonications sites, except for left 8aD, which informed both experiments). For both animals, BBB disruption was reported by *in vivo* GBCA-enhanced MRI and *ex vivo* Evans blue staining. Tissue damage was assessed using H & E staining.

**Figure 3.**
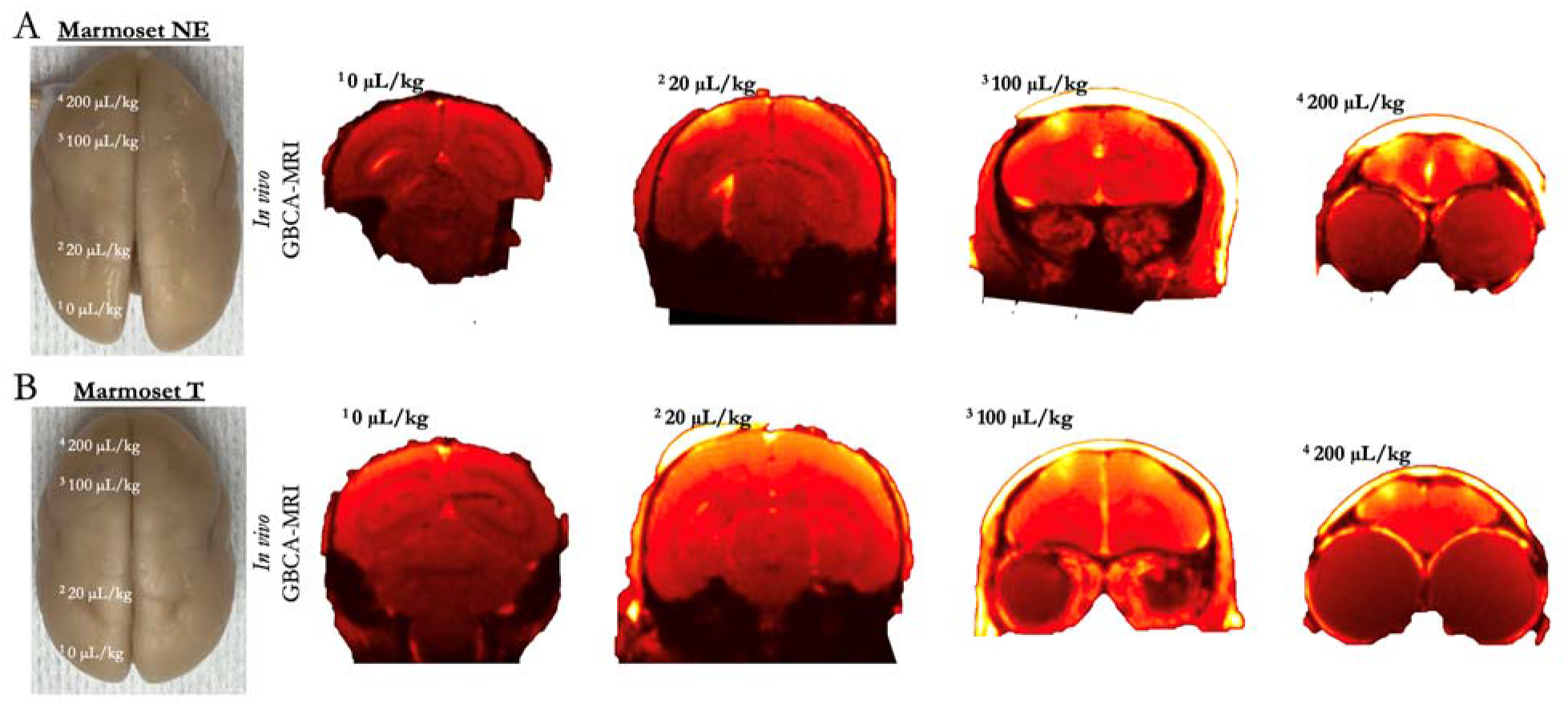
Minimum microbubble dosage for reliable blood-brain barrier disruption. (A) shows the resultant disruption (extravasation of Evans blue, left photo; and GBCA using *in vivo* MRI, right) of the BBB at the site of four sonications, with varied microbubble dosages in Marmoset NE. (B) shows the same experiment, but in marmoset T.

#### Microbubble clearance

Marmosets T and M received four sonications (parietal areas PFG right, PE right, PE left, and PFG left; albeit slightly different loci, see Figure 4 for monkey-wise locations) to determine the ability to open the BBB across multiple sites after a single bolus injection of microbubbles. Figure 4 shows sonication locations after 30, 60, 90, 120, 240, and 480 seconds after a bolus injection of 200 μL/kg of microbubbles. For all sites, a 1.46 MHz transducer was used following parameters remaining constant: acoustic pressure = 2.2 MPa, burst duration = 20 ms, burst period = 1,000 ms, number of bursts = 60. BBB disruption was reported by *in vivo* GBCA-enhanced MRI (note that the MRI sequence varied for Marmosets T and M, but both clearly showed T1-weighted GBCA contrast) and *ex vivo* Evans blue staining. Tissue damage was assessed using H & E staining. Frequency spectra from acoustic emissions were generated using a fast Fourier transform and averaged across pulses (Supplementary Figure 1).

**Figure 4.**
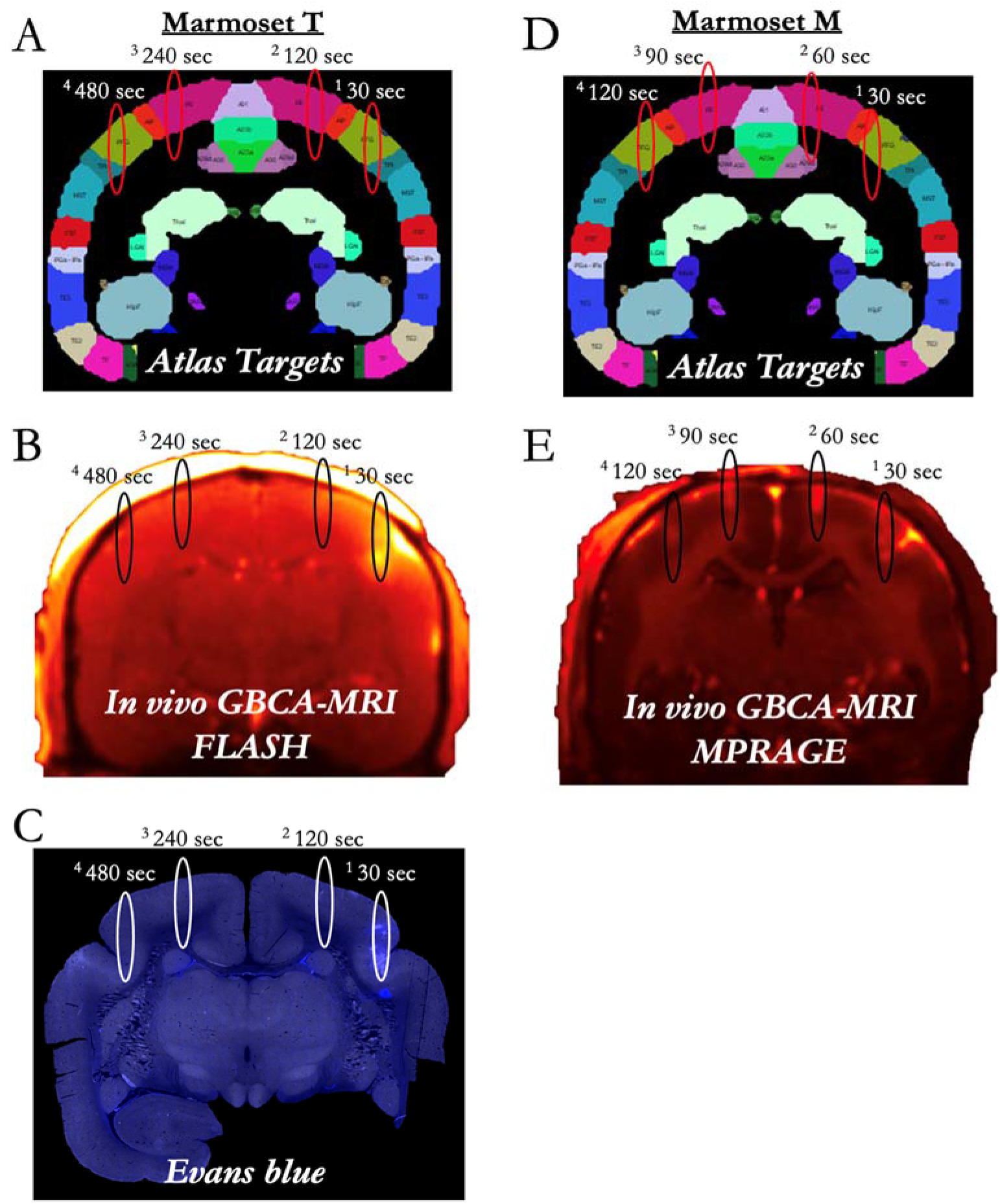
Microbubble clearance. (A) shows sonication target locations for Marmoset T at 30 – 480 seconds after a single bolus injection of 200 μL/kg of microbubbles, reported by GBCA MRI in (B) and Evans blue in (C). (D) shows similar target locations for Marmoset M, but at 30 – 120 seconds after a single bolus injection of microbubbles at 200 μL/kg, with (E) demonstrating GBCA contrast enhancement due to successful BBB disruption. Note that the MRI sequence varied for Marmosets T and M due to Marmoset M participating in other experiments, but both clearly showed T1-weighted GBCA contrast. No Evans blue was injected in Marmoset M.

#### BBB disruption as a function of skull angle

Marmoset SP received two sonications (left: 6DC; right: area 6VA). Unlike the previous experiments, the sonications were not delivered symmetrically – the right hemisphere sonications were translated lateral from the left hemisphere sonications to additionally vary skull angle along the medial-lateral axis (Figure 5 for locations). Both sonications used the following parameters with a 1.46 MHz transducer: acoustic pressure = 3.2 MPa, burst duration = 30 ms, burst period = 1,000 ms, number of bursts = 90, microbubble dosage = 400 μL/kg. A high-resolution computed tomography (CT) image was acquired to calculate skull angle and coregistered to template space^21^. BBB disruption size was calculated with the FIJI software package^28^ based on the Evans blue staining microscopy images (Figure 5E).

**Figure 5.**
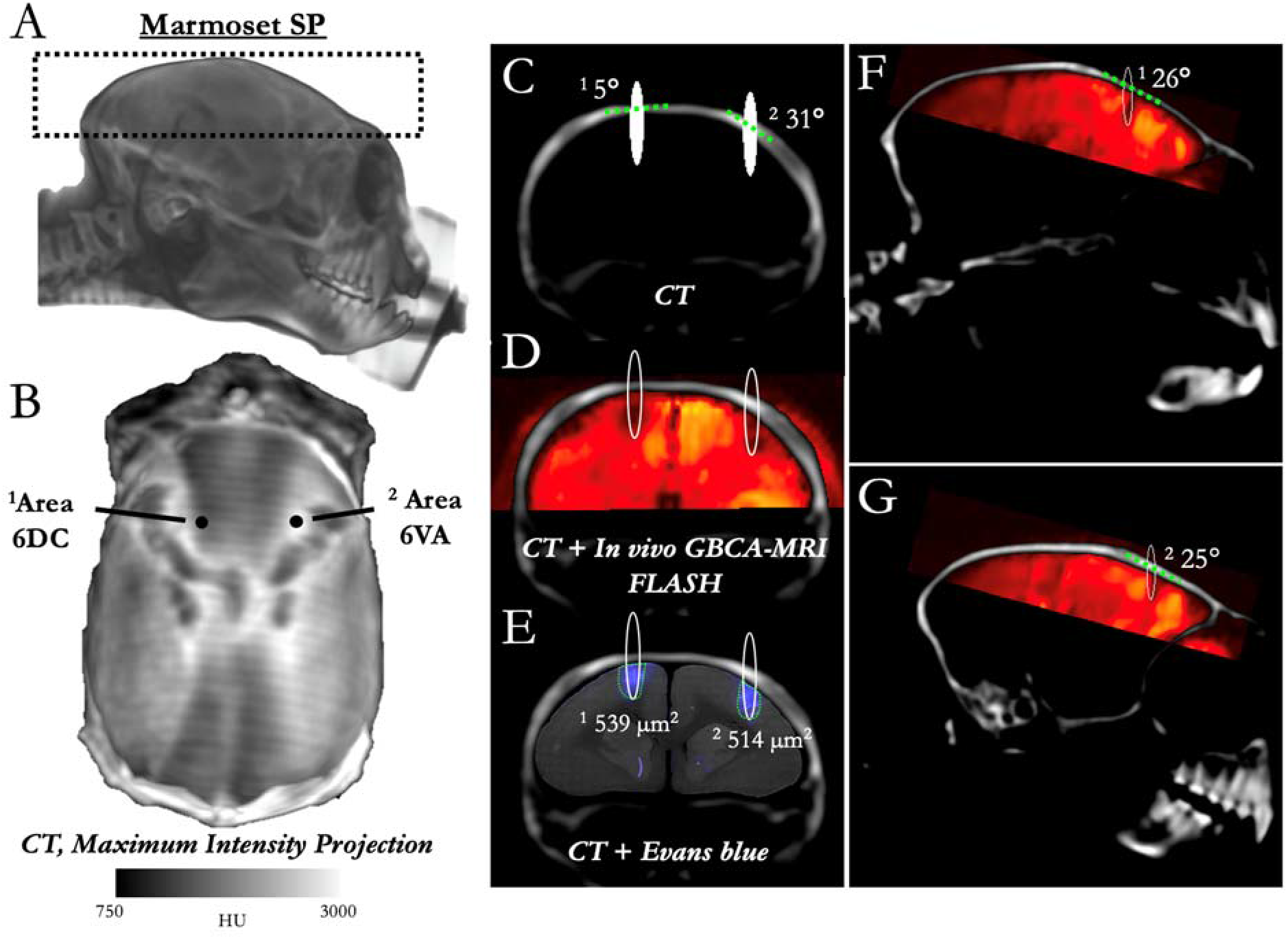
BBB disruption across varied skull thickness and angle. (A) shows a 3-dimensional rendering of marmoset SP’s CT, with the black dashed lines showing the window for the maximum intensity projection (MIP) image shown in (B). (B) shows the distribution of density (as indexed by Hounsfield Units (HU)) across marmoset SP’s skull with reference to sonication locations 1 and 2 – other than location, sonication locations 1 and 2 share the same parameters and microbubble dosage. (C) shows a coronal slice of marmoset SP’s skull with the sonication location overlaid (FWHM of the acoustic pressure distribution at the focus) and the skull angle (dashed green line tangent to skull). (D) shows the resultant opening with *in vivo* GBCA-MRI aligned to the same animal’s CT. (E) demonstrates that despite the varied thickness and skull angle, the BBB opening size was minimally affected, with Evans blue staining (microscopy image aligned to CT) showing an opening volume of sonication site 2 to be 95 % of that of sonication site 1. D & G show the same images as (D), but in sagittal slices, with the green dashed line showing the skull angle in that plane.

#### BBB disruption as a function of burst duration and number of bursts

Marmosets SK received 8 total sonications (area 8a, 4ab, MIP, and V2 bilaterally). In the left hemisphere, duty cycle was varied from 5 ms to 20 ms (Figure 6 for duty cycle by location), with the following parameters held constant with a 1.46 MHz transducer: acoustic pressure = 3.2 MPa, burst period = 1,000 ms, number of bursts = 60, microbubble dosage = 400 μL/kg. In the right hemisphere, the number of bursts was varied from 5 to 40 bursts (Figure 6 for number of bursts by location), with the following parameters held constant with a 1.46 MHz transducer: acoustic pressure = 3.2 MPa, burst duration = 20 ms, burst period = 1,000 ms, microbubble dosage = 400 μL/kg. Marmoset M received 6 total sonications with the same 1.46 MHz transducer, but with reduced parameters and microbubble dose than marmoset SK: acoustic pressure = 2.2 MPa, burst period = 1,000 ms, microbubble dosage = 200 μL/kg. In the left hemisphere (Figure 6 for locations), burst duration was varied from 5 to 20 ms; in the right hemisphere, the number of bursts was varied from 5 to 20 bursts (note, again, that the MRI sequence varied for Marmosets SK and M, but both clearly showed T1-weighted GBCA contrast).

**Figure 6.**
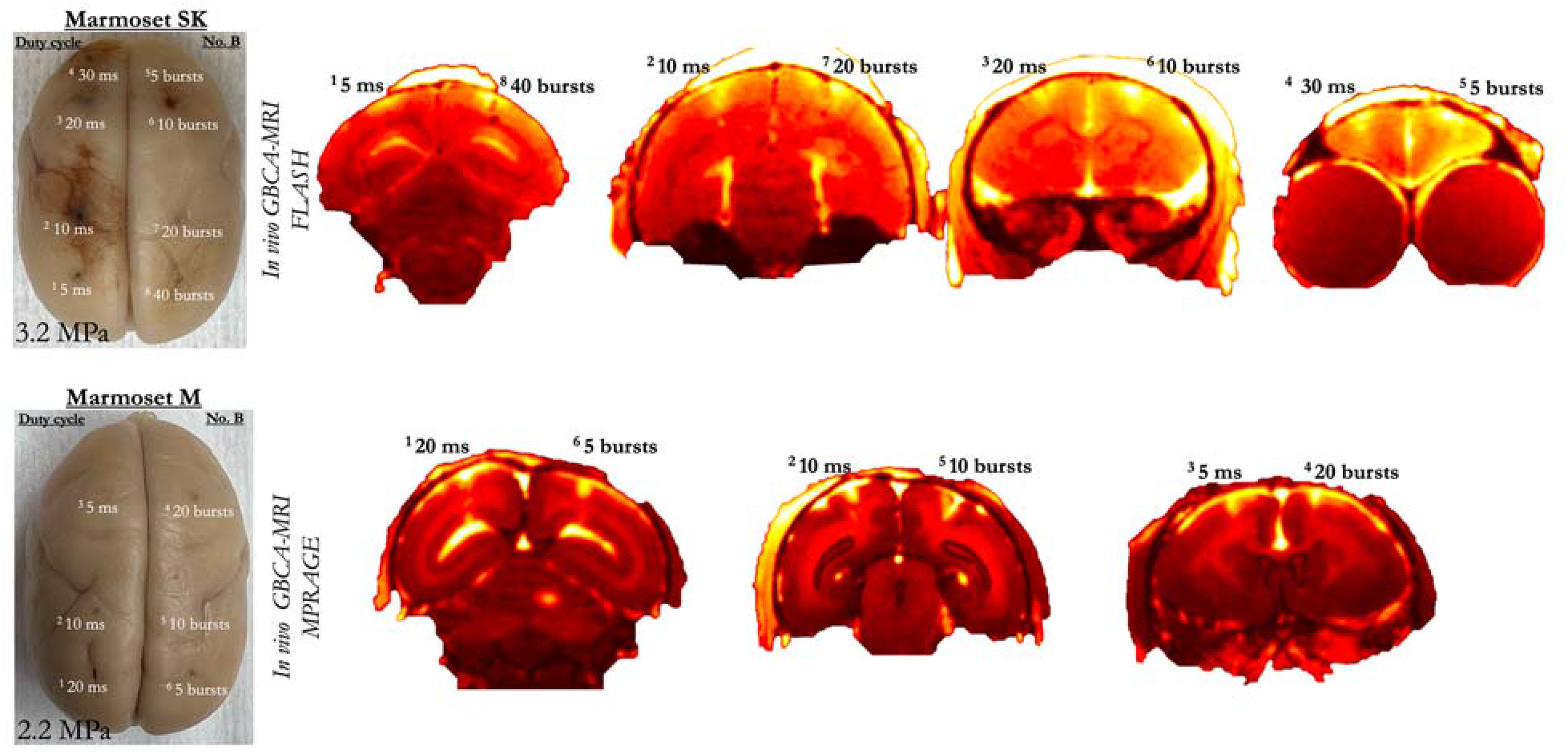
Blood-brain barrier disruption as a function of burst duration and number of bursts. (A) shows the resultant disruption (extravasation of Evans blue, left photo; and GBCA using *in vivo* MRI, right) of the BBB at the site of eight sonications, with varied duty cycle and number of bursts in Marmoset SK at 3.2 MPa. (B) shows the six sonications in Marmoset M at a lower acoustic pressure of 2.2 MPa. Note that the MRI sequence varied for Marmosets SK and M due to marmoset M participating in other experiments, but both clearly showed T1-weighted GBCA contrast.

#### BBB disruption duration

Marmoset G received a single sonication (area 8a, right hemisphere) with a 1.46 MHz transducer and the following parameters: acoustic pressure = 1.8 MPa, burst duration = 20 ms, burst period = 1,000 ms, number of bursts = 60, microbubble dosage = 100 μL/kg. BBB disruption was reported by *in vivo* GBCA-enhanced MRI. To determine the duration that the BBB remained open, Marmoset G was administered three bolus injections of a GBCA at 2-, 5-, and 8-hours post-sonication. MPRAGE anatomical images (sequence detailed below) were collected hourly to determine the changes in the size and intensity of the GBCA passing the BBB into parenchyma. BBB disruption was also reported via Evans blue, injected immediately after the last sonication. Tissue damage was assessed using H & E staining, accompanied by IBa1, NeuN, and DAPI. Marmoset M also received a single sonication in area 8a (2.2 MPa, burst duration = 20 ms, burst period = 1,000 ms, number of bursts = 60, microbubble dosage = 200 μL/kg), but the GBCA was administered at 2 hours, then a much longer period of 2 weeks after sonication. Given the survival time necessary for the experiment (2 weeks), Marmoset M did not receive Evans blue.

### Magnetic resonance imaging

#### In vivo MRI, CT

All neuroimaging (MRI & CT) took place at the University of Pittsburgh Brain Institute. For MRI, a 9.4 T 30 cm horizontal bore scanner (Bruker BioSpin Corp, Billerica, MA) was used, equipped with a Bruker BioSpec Avance Neo console and the software package Paravision-360 (version 3.2; Bruker BioSpin Corp, Billerica, MA), and a custom high performance 17 cm gradient coil (Resonance Research Inc, Billerica, MA) performing at 450 mT/m gradient strength. To detect BBB disruption *in vivo*, MRI was acquired on Marmosets B, SG, NE, T, M, SK, SP and G in concert with an intravenous contrast agent (Marmoset B: manganese chloride; all others: gadolinium) that was injected after the sonications (within 1 minute of last sonications, except Marmoset E to determine BBB disruption duration) and before the MRI (MRI started within 20 minutes of transferring the animal from the FUS including scanner preparations involving localization and magnetic field shimming). Radiofrequency transmission was accomplished with a custom 135 mm inner diameter coil and a custom in house 8-channel phased-array marmoset-specific coil was used for radiofrequency receiving. Marmosets were imaged in the sphinx position, with a custom 3D printed helmet for head fixation and anesthesia mask for inhalant isoflurane delivery. A T1-weighted fast low angle shot (FLASH) sequence was employed to detect the resultant shortening of T1 relaxation times from the contrast agents entering the parenchyma via the BBB disruption. Three scans (later averaged) were acquired for each animal with the following parameters: TR = 25 ms, TE = 8 ms, field of view = 35 x 35 x 26 mm, matrix size = 117 x 117 x 87, voxel size = 0.299 x 0.299 x 0.299 mm, bandwidth = 200 kHz, flip angle = 25 degrees, total scan time = 9 minutes, 2 seconds. For marmosets G & M, a magnetization prepared – rapid gradient echo (MPRAGE) sequence was used in lieu of the FLASH sequence because of the longitudinally-additive effects of systemic GBCA on the dynamic signal contrast. With the MPRAGE sequence, this additive effect is reduced with the additional inversion pulse. The MPRAGE sequence was acquired with the following parameters: TR = 6,000 ms, TE = 3.42 ms, field of view = 42 x 35 x 25 mm, matrix size = 168 x 140 x 100, voxel size = 0.250 x 0.250 x 0.250 mm, bandwidth = 50 kHz, flip angle = 14 degrees, total scan time = 20 minutes, 6 seconds.

To accurately quantify skull angle relative to the ultrasonic transducer, CT was acquired on Marmoset SP at the University of Pittsburgh Brain Institute on a small animal CT (Si78; Bruker BioSpin GmbH, Ettlingen, Germany) equipped with the software package Paravision-360 (version 3.1; Bruker BioSpin Corp, Billerica, MA). Using a Low Dose 1 mm aluminum filter, a 200 x 200 x 200 μm (field of view = 79.6 x 81.1 mm) CT was acquired using a “step and shoot” method (0.6-degree gantry step) and reconstructed using the filtered back projection algorithm. The MRI and sliced microscopy images were then aligned using the 2D to 3D registration provided in DSIstudio^29^.

### Histology and immunohistochemistry

After *in vivo* MRI acquisition, all nine marmosets were euthanized with pentobarbital sodium and phenytoin sodium solution (100 mg/kg) for histological examination. Transcardial perfusion was performed with 4 % paraformaldehyde. The brains were removed, postfixed, and cryoprotected in 30 % sucrose for 3-5 days. Marmoset brains were sectioned coronally at 30 μm using a cryostat (Leica CM1950, Deer Park, IL, USA) and stored in cryoprotectant solution with 15 % glycerol and 15 % ethylene glycol at −20 °C until further use. For Evans blue fluorescence, sections were mounted onto Superfrost slides (Fisher Scientific) and visualized under an AxioImager M2 epifluorescence microscope (Car Zeiss, White Plains, NY, USA). For Hematoxylin and Eosin (H & E) staining, sections were mounted onto a Superfrost slide and stained with an H & E stain kit (#3502 Vector Laboratories, Newark, CA, USA) by following procedures suggested by the manufacturer. Images were captured using an AxioImager M2 microscope (Carl Zeiss). For immunofluorescence staining, floating sections were permeabilized in blocking buffer (2 % donkey serum and 0.2 % Triton X-100 in PBS) at room temperature for 1 h with gentle shaking, followed by overnight incubation with primary antibodies, Iba1 (Ionized calcium binding adaptor molecule 1 - 1:500; #019-19741 Wako Chemicals) and NeuN (Fox-3 - 1:500; #MAB377 MilliporeSigma) at 4 °C. After PBS wash, sections incubated with fluorescent secondary antibodies, Alexa Fluor 488-conjugated donkey anti-rabbit IgG (Invitrogen) and Alexa Fluor 647-conjugated donkey anti-mouse IgG (Invitrogen). Sections were counterstained with DAPI (4’,6-Diamidine-2’-phenylindole dihydrochloride - Invitrogen) for staining nuclei. Images were acquired using a LSM900 confocal microscope (Carl Zeiss) using 10× objective at 1024 × 1024 pixel resolution with z-step size of 1 μm thickness.

## Results

### BBB disruption as a function of center frequency

Marmoset B received two sonications in parietal cortex to determine the relative difference in BBB disruption size as a function of transducer center frequency. With the goal of minimizing the size of disruption, both transducers (515 kHz, left hemisphere; 1.46 MHz, right hemisphere) were focused on the edge of the cortical ribbon (Figure 1C) – consequently, only about half of the ultrasonic beam in the axial orientation was focused on brain tissue. As evidenced by the Evans blue staining and MnCl2-enhanced MRI (Figure 1C), both sonications were close in volume to the full width at half maximum of the acoustic pressure distribution at the focus (with 515 kHz at 188 mm^3^ and 1.46 MHz at 10 mm^3^). With the aim of subsequent experiments to demonstrate focal BBB disruptions, only the higher-frequency 1.46 MHz transducer was used. As reliably demonstrated across the experiments described below, the method of centering the focus of the 1.46 MHz transducer at the top of the cortex allowed for disruptions on the order of 1 mm radially, and 2.5 mm axially. Indeed, the disruption size varied slightly as a function of acoustic pressure, microbubble dosage, burst duration, and number of bursts, but at the “safe” parameters demonstrated below, the 1.46 MHz transducer allowed for cortical disruptions within the bounds of most cortical cytoarchitectonic regions of interest (e.g., fit within the medial-lateral extent of area MIP).

### BBB disruption as a function of acoustic pressure

Marmosets SG, NE, and T each received five or eight cortical sonications (area 8a, 4ab, MIP, and V2) with a 1.46 MHz transducer. Figure 2 shows the sonication locations and accompanying acoustic pressures. Based on the initial experiments in marmoset SG demonstrating that acoustic pressures of 1.8 MPa – 3.2 MPa (and a high 400 μL/kg microbubble dose) allowed for reliable BBB disruption, marmosets NE and T were sonicated with lower pressures of 0.6 MPa – 2.2 MPa and a lower 200 μL/kg microbubble dose to determine the minimum pressures at which perturbing the BBB allows for extravasation of GBCA and Evans blue stain. As shown in Figure 2, 1 MPa perturbed the BBB in marmoset NE, but not marmoset T. 1.4 MPa successfully perturbed the BBB in both marmoset NE and T, but there was a clear superficial cortical bias of the distribution of BBB disruption closer to the center of the focus of the acoustic beam. At 1.8 MPa, marmoset NE showed a clear perturbation at a size approximating volume to the full width at half maximum of the acoustic pressure distribution at the focus (2.5 mm axially by 1 mm radially), but less so in marmoset T. At 2.2 MPa marmoset SG, NE, and T showed a consistent disruption (note that the GBCA agent dosage for marmoset SG was lower at 0.1 ml/kg, but see photograph of Evans blue staining in Figure 2 at 2.2 MPa).

### Minimum microbubble dosage to disrupt the BBB

The right hemispheres of marmosets NE and T were used to determine the minimum acoustic pressure (Figure 2), while the left hemisphere was used to demonstrate the minimum microbubble dosage necessary to perturb the BBB (Figure 3). Each animal was sonicated four times (area 8a, 4ab, MIP, and V2), with the lowest microbubble dosage used for the most posterior sites from 0 μL/kg (V2), then increasing to 20 μL/kg (MIP), 100 μL/kg (4ab), and 200 μL/kg (8a) at the anterior sites. Though the parameters (and timing) were otherwise the same for marmosets NE and T, 20 μL/kg perturbed the BBB in marmoset NE, but not fully in marmoset T. 100 μL/kg successful perturbed the BBB in both animals, as did 200 μL/kg. Thus, with our apparatus, we demonstrate minimum microbubble dosage for single-bolus tail vein injections in marmosets to be in the range of 20 – 100 μL/kg at 1.46 MHz.

### Microbubble clearance

With a total of 12 sonications (8 of which are described in the previous two sections), Marmoset T also received four sonications (parietal areas PFG right, PE right, PE left, and PFG left) to determine the ability to disrupt the BBB across multiple sites after a single bolus injection of microbubbles. Figure 4 shows sonication locations after 30, 120, 240, and 480 seconds after a bolus injection of 200 μL/kg of microbubbles. At 2.2 MPa, only the sonication 30 seconds after microbubble injection showed perturbation which allowed for clear extravasation of GBCA and Evans blue stain at the approximate volume at FWHM of the acoustic pressure distribution at the focus, as visualized by the ellipsoids in Figure 4A, B, & C. Marmoset M shared the same sonication parameters and was used to determine the ability to disrupt the BBB at 30, 60, 90, and 120 seconds (between the first two points of Marmoset T) after a single bolus injection. From this data, the effective clearance time of a 200 μL/kg dose of commercially available microbubbles (Definity) appears to be less than two minutes in the marmoset brain – we expect that there is some variation in effective concertation of intravascular microbubbles as a function of the animal’s physiological state (e.g., heart rate, kidney function). The dose here, however, was relatively high at 200 μL/kg, so the availability of bubbles in circulation could not be much higher without leading to damage (see Figure 8). Across the (other) experiments presented here, we injected prior to each sonication rather than using a single bolus or infusion for multiple sonications, with at least 5 minutes between injections. Likely due to variation in tissue (e.g., muscle, fat) between the sonicated site and the hydrophone, we did not find the Fourier spectrum of the monitored cavitation emissions to be a particularly reliable index of BBB perturbation (with ground truth from the Evans blue staining microscopy). However, this experiment allowed for the opportunity of comparing spectra from successful disruptions to unsuccessful ones with identical parameters, only varying available microbubbles in circulation. From these four sonications (marmoset T), broadband noise and an increase in subharmonic amplitude around half of the transducer frequency (.73 MHz) appeared in relation to cavitation corresponding to disruption at 30 and 120 seconds post microbubble injection, but only the 30 second post-microbubble injection sonication showed successful BBB opening. This opening was also accompanied by an increase in amplitude at the second harmonic (Supplementary Figure 1).

**Figure 7.**
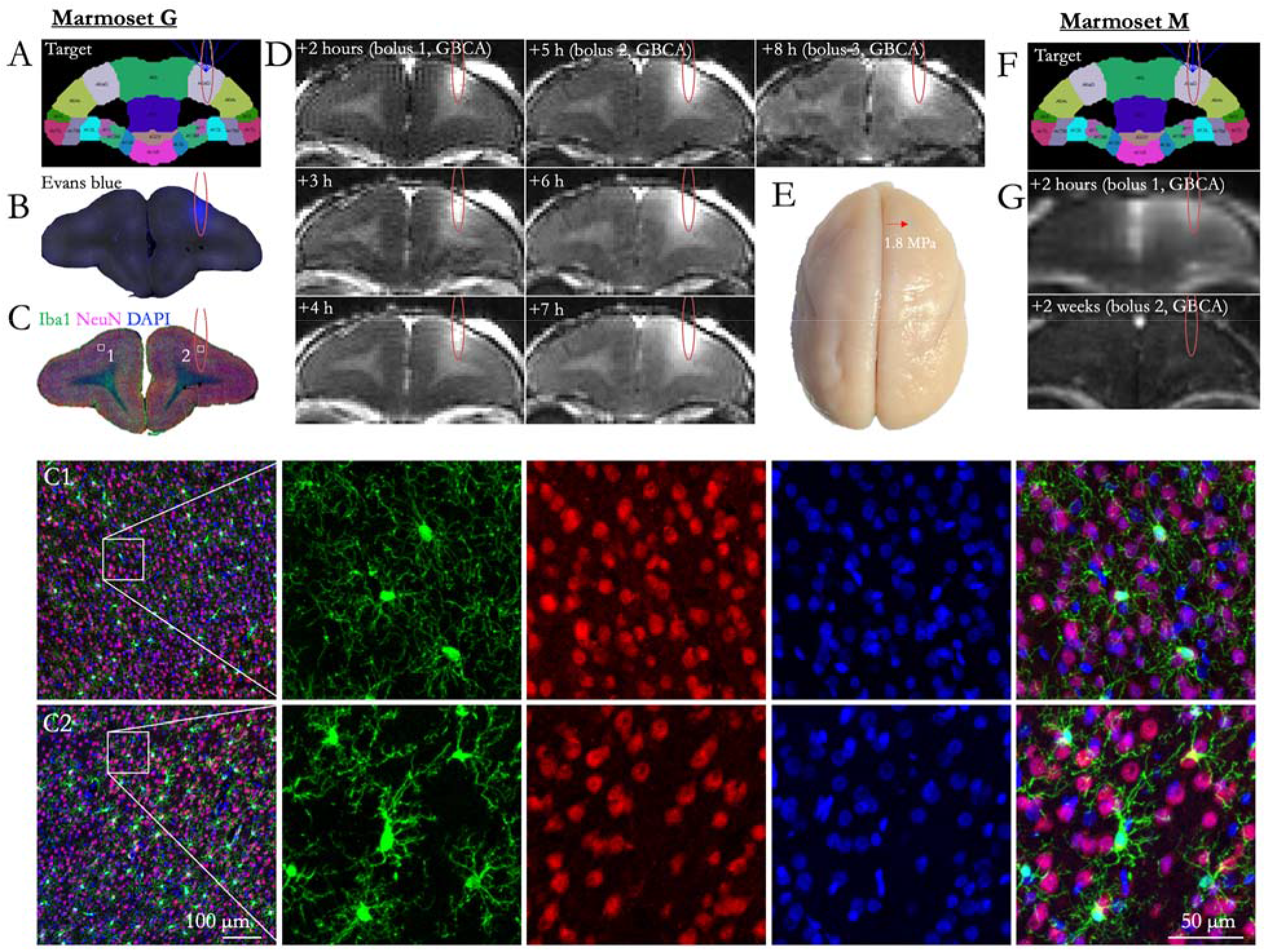
Blood-brain barrier opening duration and immunohistochemistry. (A) Shows the target for a single sonication for Marmoset G, area 8a. (B) shows the extravasation of Evans blue as the result of this sonication. (C) shows immunostaining at the sonicated slice, with C1 showing the left (not sonicated) hemisphere and C2 showing right hemisphere within the bounds of the sonicated area. Compared to left hemisphere, a marked increase in Iba1+ cells was clearly evident in right hemisphere. Iba1+ cells exhibit larger cell bodies and thick processes in right hemisphere, whereas Iba1+ cells show much more numerous and finer processes in left hemisphere. (D) shows *in vivo* GBCA MRI, with boluses of GBCA injected at 2, 5, and 8 hours after sonication. As shown in the hours following the bolus injections (hours 3 and 4, also hours 6 and 7), the size of the GBCA remained the same (i.e., minimal diffusion), but increased with subsequent bolus injections at hours 5 and 8, suggesting the BBB was still open at 8 hours post sonication. (E) shows a photograph of Marmoset G’s perfused brain, with the sonication site marked by a red arrow, indicating no evidence of hemorrhage at this site. (F) shows the target for a single sonication in Marmoset M, area 8a. (G) shows the extravasation of GBCA, injected as a bolus two hours after sonication, then a second bolus injection again after two weeks. After two weeks, the BBB was no longer permeable to the GBCA.

**Figure 8.**
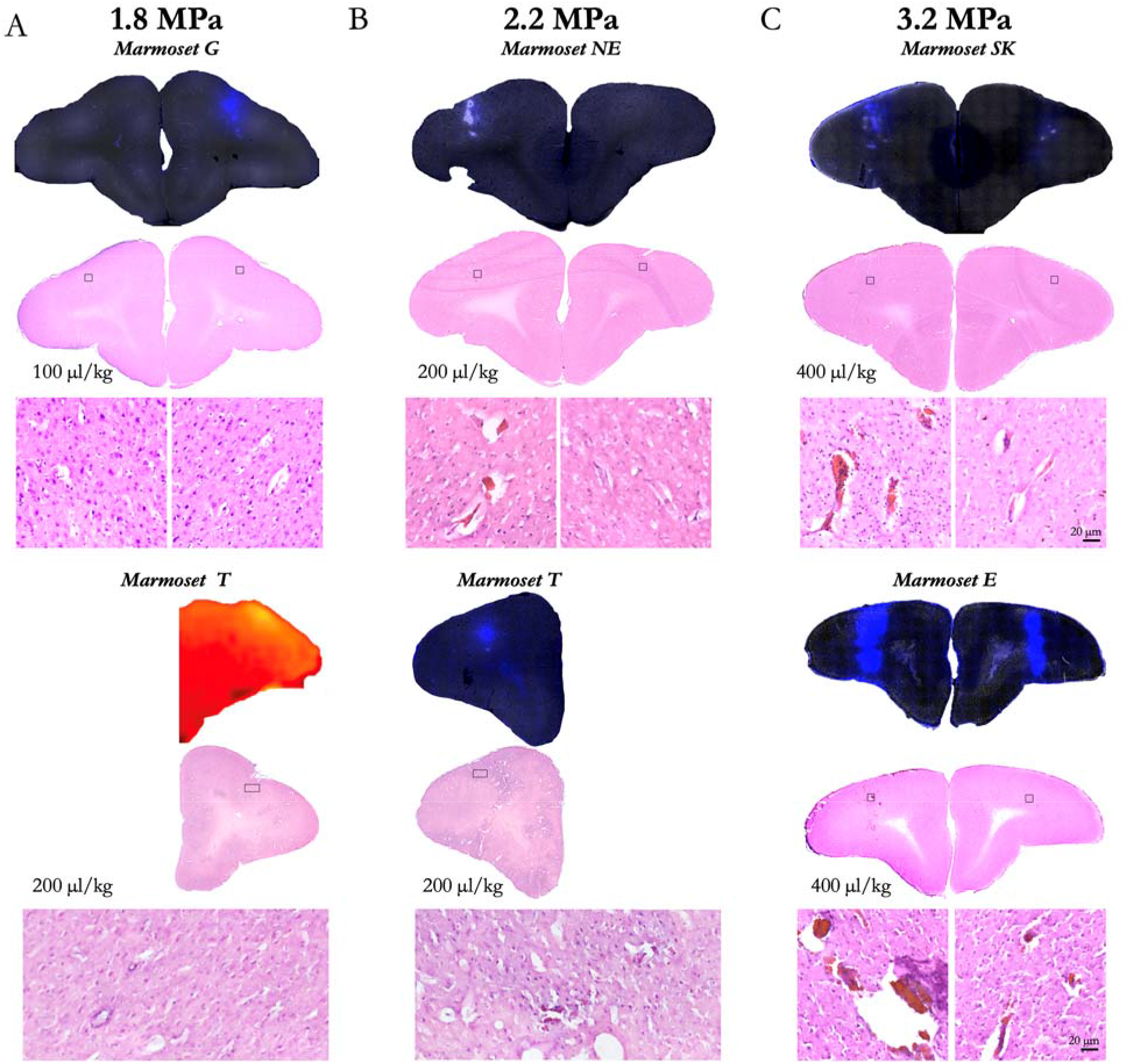
Cortical damage as a function of acoustic pressure and microbubble dosage. (A) show sonications at 1.8 MPa, (B) at 2.2 MPa, and (C) at 3.2 MPa. For each acoustic pressure (columns) and microbubble dose (rows), Evans blue microscopy images or GBCA images are shown colocalize the site of sonication with the H & E stained slice of interest. Black boxes overlaying the H & E images show the location of the zoomed image below each slice of interest.

### BBB disruption as a function of skull angle

With known effects of acoustic reflection mediated by skull angle and thickness^30^, we sought to directly test these effects by sonicating at 1.46 MHz across varied skull angles of the marmoset head. As shown in the apparatus rendering in Figure 1, our transducer was always parallel with the stereotactic plane (i.e., the Z plane corresponding to the center of the ear bars and bottom of the eye bars) and thus was fixed with reference to the varying skull angle. In addition to *in vivo* GBCA-enhanced MRI, high-resolution (50 μm isotropic) CT images were acquired in marmoset SP to isolate the skull and accurately measure angles tangent to the skull surface (tangent to the point at the intersection of the axial center of the sonication ellipsoid). At high pressure (3.2 MPa, 400 μL/kg microbubble dose), the size of the BBB disruption matched the FWHM of the expected sonication distribution, indicating the sonications planar to (above the) marmoset skull were not adversely affected by skull angle. Accordingly, the thin marmoset skull (~1 mm or less) seems to be amenable to focused ultrasound compared with other species. It is also worth noting that although the acoustic pressure and microbubble dose were high for this experiment, data from Figures 2, 3, 6 or 7 also demonstrate that skull angle seems to have a minimal effect at lower acoustic pressures and microbubble dosages. Because of the focus of our transducer (focal length = 24.5 mm), we have yet to establish these effects deeper within the brain. With the transducer used here, we could only effectively center the acoustic beam 11 mm from the top of the marmoset head, limiting the ability to establish such effects.

### BBB disruption as a function of burst duration and number of bursts

Marmosets SK received 8 sonications and Marmoset M received 6 sonications to determine the effect burst duration and number of bursts on the extent of BBB disruption – and, although unintentional, the extent and nature of damage at high acoustic pressure for Marmoset SK. Indeed, for marmoset SK, with such a high pressure and microbubble dosage, all the tested duty cycles (5, 10, 20, and 30 ms) and number of bursts (5, 10, 20, 40) disrupted the BBB in marmosets SK. These data, however, are particularly useful for demonstrating the extent of damage that can occur at the extremes of acoustic pressure, burst durations, and at high microbubble dosages (400 μL/kg), as shown in Supplementary Figure 2. For Marmoset M, however, the pressure (2.2 MPa) and microbubble dosage (200 μL/kg) were closer to the boundary of safe sonication parameters (Figure 8). Of the tested parameters, as low as 5 bursts (20 ms burst duration) or 5 ms burst duration (60 bursts), we found that we were likely above the minimum to disrupt the BBB. These data suggest that the disruption likely occurs with the first few bursts, when the circulating microbubble concertation is highest. This bodes particularly well for applications in which very fast sonications are advantageous, such as those delivered while the animal is awake and performing a task.

### BBB disruption duration

Marmoset G received a single sonication (area 8a, right hemisphere) with a 1.46 MHz transducer and the following parameters: acoustic pressure = 1.8 MPa, burst duration = 20 ms, burst period = 1,000 ms, number of bursts = 60, microbubble dosage = 100 μL/kg. As shown in Figure 8, BBB disruption was reported by *in vivo* GBCA-enhanced MRI and Evan’s blue staining. From the previous experiments and the results shown in Figure 8, we found these parameters to be both safe and effective at opening the BBB. In addition to demonstrating the safety and reliability of opening with the aforementioned parameters, we sought to determine the length of time that BBB remained open. As this was a scheduled terminal experiment (with Evans blue already injected, precluding recovery of the animal), we were not able to determine the upper limit BBB disruption duration. Rather, our last point of imaging was 8 hours after injection. As shown in Figure 8, the BBB was clearly still open at 8 hours post-sonication as indicated by increased intensity and distribution resulting from the GBCA and at the start of the 8-hour post-sonication scan. Marmoset M, however, was not injected with Evans blue and was injected with GBCA 2 weeks after sonication – the BBB no longer allowed passage of a sufficient volume of GBCA to be detected. Note that the images from Figure 6 were acquired at the same time as Marmoset M’s image in Figure 7 (demonstrating that the GBCA injection was successful), with extravasation of GBCA at the sites sonicated 2 weeks after the first 8a sonication.

#### Histology and immunohistochemistry

To determine efficiency and safety of BBB disruption according to changes in acoustic pressure and microbubble dosage, both Evans blue extravasation and histological tissue damage were confirmed in response to acoustic pressure of 1.8, 2.2, and 3.2 MPa with 100, 200, or 400 μL/kg microbubble dosage. As shown in Figure 8, Evans blue extravasation was observed for acoustic pressure of 1.8 MPa and higher acoustic pressure (2.2 and 3.2 MPa). Acoustic pressure of 3.2 MPa with 400 μL/kg microbubble dosage produced severe tissue damage and large clusters of erythrocyte extravasation in marmoset SK, whereas an acoustic pressure of 1.8 MPa facilitated BBB disruption in the absence of apparent tissue damage or microhemorrhages in marmoset G and T. Moreover, there was no apparent neuronal damage identifiable in this region when examining the NeuN immunostaining (Figure 7). Ionized calcium-binding adapter molecule 1 (Iba1) expression is upregulated upon microglia activation. Therefore, we used immunostaining of this microglial marker as measures of immune response. Compared to contralateral hemisphere, a marked increase in Iba1^+^ microglia was clearly evident in ipsilateral hemisphere (Figure 7). Iba1^+^ cells exhibited larger cell bodies and thick processes in ipsilateral hemisphere, whereas Iba1^+^ cells showed much more numerous and finer processes in the contralateral hemisphere.

## Discussion

In this study, we sought to establish the parameters to focally disrupt the BBB across a cohort of marmoset monkeys. By integrating our MRI-based marmoset atlases^21,22^ with a motorized stereotactic positioning system (RK-50 Marmoset; FUS Instruments Incorporated, Toronto, ON, Canada, Figure 1) we were able to focally sonicate sites across the dorsal surface of the marmoset cortex with high accuracy (Figures 1-8) by way of spherically focused single-element transducers. Of the two transducers tested – at 515 kHz and 1.46 MHz – we found that the higher frequency 1.46 MHz transducer (with a 24.5 mm focal length) allowed for disruptions that could be limited to the ~2.5 – 3 mm cortical thickness of the marmoset cortex. We optimized parameters (minimum acoustic pressure, minimum microbubble dosage, burst duration, number of bursts) from which BBB disruption occurred without hemorrhage or edema and to the extent that Evans blue and/or GBCA extravasation occurred. From these experiments, we establish parameters for safe BBB disruption (as reported by H & E staining and immunohistochemistry) in the marmoset at 1.46 MHz. At these parameters, we found that spatial bounding of Evans blue staining and GBCA reporting (Figure 7) were similar, such that the determination of a BBB disruption can be conducted *in vivo* with MRI-based contrast agents. Taken together, the experiments described here provide an account from which *in vivo*, minimally invasive substance delivery experiments can be designed around.

As with the rodent brain, marmosets have a lissencephalic cortex, making this species an ideal candidate for tFUS-based BBB disruption, allowing for sonications along the columnar organization of the cortex, unencumbered by cortical folds present in most other primate species. As we show here, however, sonication parameters from rodent species (e.g., rats, mice)^8^ cannot be simply ported for use in the marmoset, nor can those used in for other larger nonhuman primate species such as macaques (with a much thicker skull, and folded cortex)^4^. Indeed, in addition to the parameters driving the transducer, the physical properties of transducers also complicate comparisons (e.g., focal length, frequency). The experiments presented here demonstrate that a 35 mm single element spherically focused 1.46 MHz transducer can be used for cortical disruptions in the marmoset with minimal deleterious effects of the marmoset head morphology (e.g., skull thickness or the presence of temporalis muscles; Figure 5), particularly when the center of focus is at or near the surface of cortex (Figures 1-8), allowing for further spatial minimization of cortical disruption. The use of a single-element transducer rather than an array of transducers simplifies the means necessary to conduct tFUS in the marmoset. We made use of a motorized positioning system and automated atlas targeting, but these experiments could also be conducted with a transducer mounted to a stereotactic manipulator arm, further simplifying the equipment needed to use focused ultrasound to sonicate the marmoset brain.

Although it should be noted that microbubble experiments (dosage, clearance time) may be parameter and transducer-dependent^31^, we found that the minimum microbubble dosage (Definity, Lantheus Medical Imaging, Billerica, MA, USA), via the tail or saphenous vein was greater than 20 μL/kg when injected as a bolus (Figure 3). To target a smaller distribution of microbubbles (with larger bubbles being more buoyant), we drew from as close to the bottom of the microbubble vial as possible (after activation and slowly inverting the vial) with a 21-gauge needle, and another 21-gauge needle to vent the vial. We also chose to use a 26-gauge catheter to avoid premature destruction of the microbubbles during injection of our saline-diluted microbubble solution. During initial experiments, we found that the use of a spring-loaded extension led to inconsistent BBB perturbation, likely due to premature bursting of the microbubbles. As such, all injections were made directly into the catheter hub. In terms of microbubble clearance time, we found it to be relatively fast (< 2 minutes) in the marmoset, even with a high dose of 200-400 μL/kg – at least to the extent that the circulating microbubble concentration resulted in BBB disruption (Figure 4). We found that the acoustic emissions indicated cavitation of the microbubbles at 30 seconds. In particular, consistent with previous reports in rats^8^ using a similar transducer and hydrophone hardware, subharmonic broadband noise was evident at 30 and 120 seconds post microbubble bolus injection, as well as an increased amplitude at the second harmonic (Supplementary Figure 1). This likely corresponded to microbubble cavitation, although it did not correspond to opening at 120 seconds post-microbubble injection. This subharmonic effect was not apparent at 240- or 480-seconds post-microbubble injection.

Our original intent to determine BBB disruption as a function of burst duration and number of bursts was not to assess damage (visible to the naked eye, Figure 5), but as shown by the H & E staining (supplementary Figure 2), the pressure used for marmoset SK was above of what can be considered safe (Figure 8). These data, however, are particularly useful for demonstrating the extent of damage that can occur at the extremes of acoustic pressure, burst durations, and at high microbubble dosages (400 μL/kg). We found that these high-energy sonications resulted in diffuse tissue damage usually accompanied by microbleeds. Of particular interest, we found that the subdural space presented hemorrhage. This area is particularly sensitive for hemorrhages due to the numerous presence of perforating arteries in a compact space and thus the microbubble concentration may have been higher.

Given the results of the H & E staining of the animals that showed reliably successful disruptions of the BBB (Figure 2 – 6), we applied what we found to be safe parameters (acoustic pressure = 1.8 MPa, burst duration = 20 ms, burst period = 1,000 ms, number of bursts = 60, microbubble dose = 100 μL/kg) to a frontal site (area 8a) in marmoset G to determine the length of time that the BBB remained open, as indexed by permeability to a bolus of GBCA at 2, 4, and 8 hours after sonication. The BBB was clearly open at 8 hours post-sonication, as indicated by increased intensity and distribution resulting from the GBCA and at the start of the 8-hour post-sonication scan (Figure 7). Although this long-duration disruption may present some risk (e.g., blood-borne bacteria) that the BBB would normally protect against circulating toxins or pathogens in the bloodstream^32^, it is an experimentally advantageous treatment window for injecting substances that may be dangerous if injected as a bolus, rather than slowly infused. The histology and immunohistochemistry in marmoset G supported the previous results that the parameters were safe, such that no readily apparent damage was observed in the H & E staining (Figure 8). There is visible microglial activation (Figure 7) due to tissue perturbation (Iba1), but not tissue damage (DAPI, NeuN, H & E) compared to a contralateral non-sonicated 8a site indicating that these are safe and reliable parameters to open the BBB for an extended period. Microglia in cortical regions showed signs of activation through increased Iba1 expression and changes in cell bodies and processes, without significant changes in cell numbers. FUS-induced BBB disruption has been shown to trigger transient glial activation^33^. Depending on the type and severity of brain injury, activated microglia as well as infiltrating macrophages can exacerbate neuroinflammation and neurodegeneration. It has been demonstrated, however, that microglia activation resolved by 15 d after FUS with no progression to a glial scar, suggesting that FUS does not cause lesion-like microgliosis^34^.

In summary, we demonstrate safe and effective disruption of the BBB in the marmoset with a spatial specificity of approximately 1 mm radially and 2.5 mm axially (in cortex) using a 1.46 MHz transducer. We were able to reliably perturb the BBB across the dorsal surface of the marmoset cortex with a minimum acoustic pressure to be between 1.8 – 2.2 MPa and a minimum microbubble dosage of 20 μL/kg via tail-vein injection. We demonstrate that these parameters (paired with 60 20 ms bursts, spaced at 1000 ms) led to the BBB being open for greater than 8 hours and did not lead to cortical tissue damage, as reported by H & E staining. Higher acoustic pressures and/or excessive microbubble dosage (> 200 μL/kg) led to tissue damage (Figure 8). The series of experiments presented here establish methods for safely, reproducibly, and focally perturbing the BBB using tFUS in the common marmoset monkey.

## Author Notes

We wish to thank Brianne Stien, Lauren Dubberly, and Dr. Julia Oluoch for animal care and preparation. This work was supported by the National Institute of Neurological Disorders and Stroke of the National Institutes of Health under Award Number R21NS125372 (D. J. S.) and by the Pennsylvania Department of Health Commonwealth Universal Research Enhancement (C.U.R.E.) Tobacco Appropriation Funds – Phase 18 (SAP 4100083102; A. C. S.). The content is solely the responsibility of the authors and does not necessarily represent the official views of the National Institutes of Health.

---

**Supplementary Figure 1.**
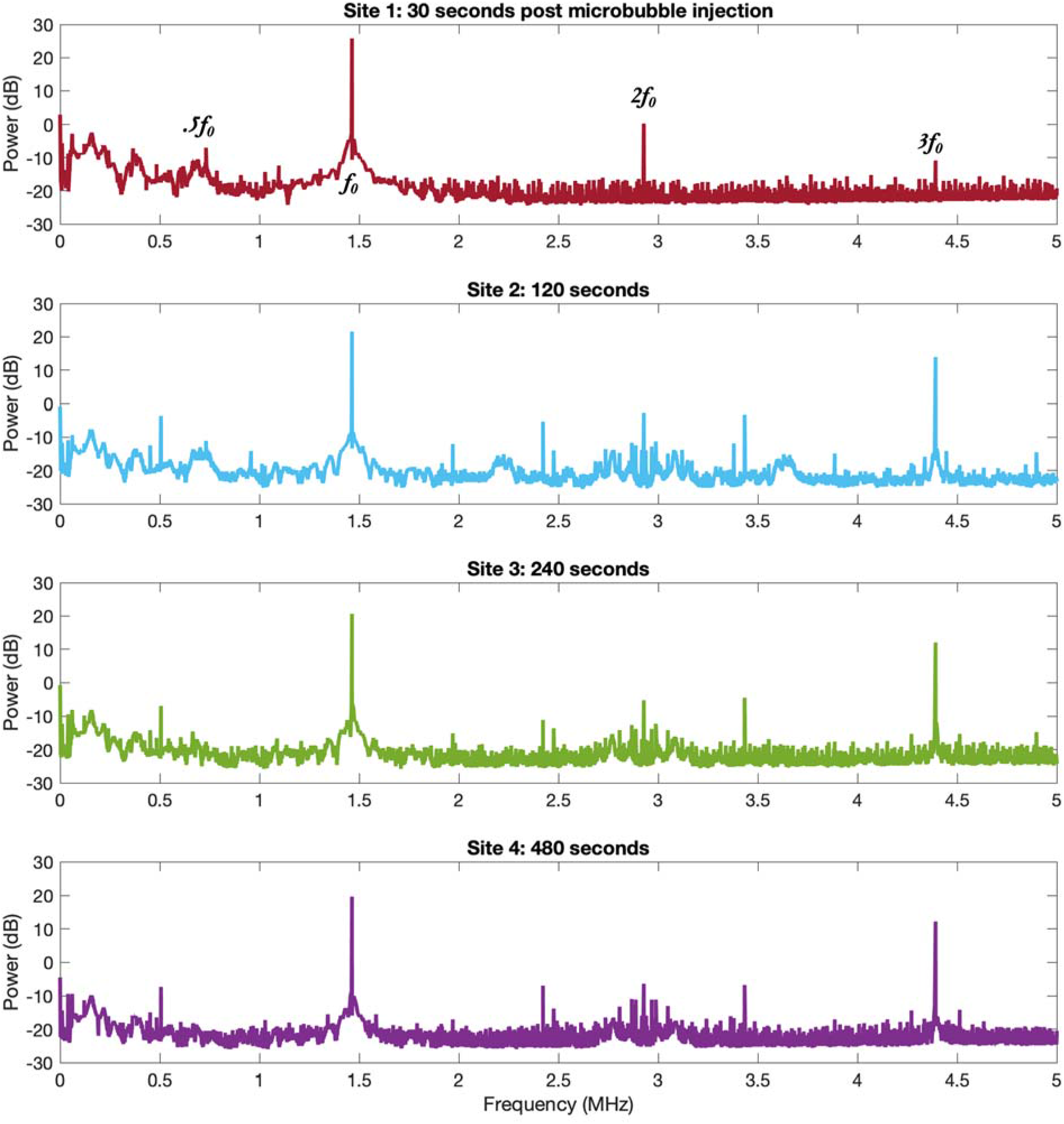
Frequency spectra from sonications in marmoset T (sites shown in Figure 4) following a single bolus microbubble injection of 200 μl/kg.

**Supplementary Figure 2.**
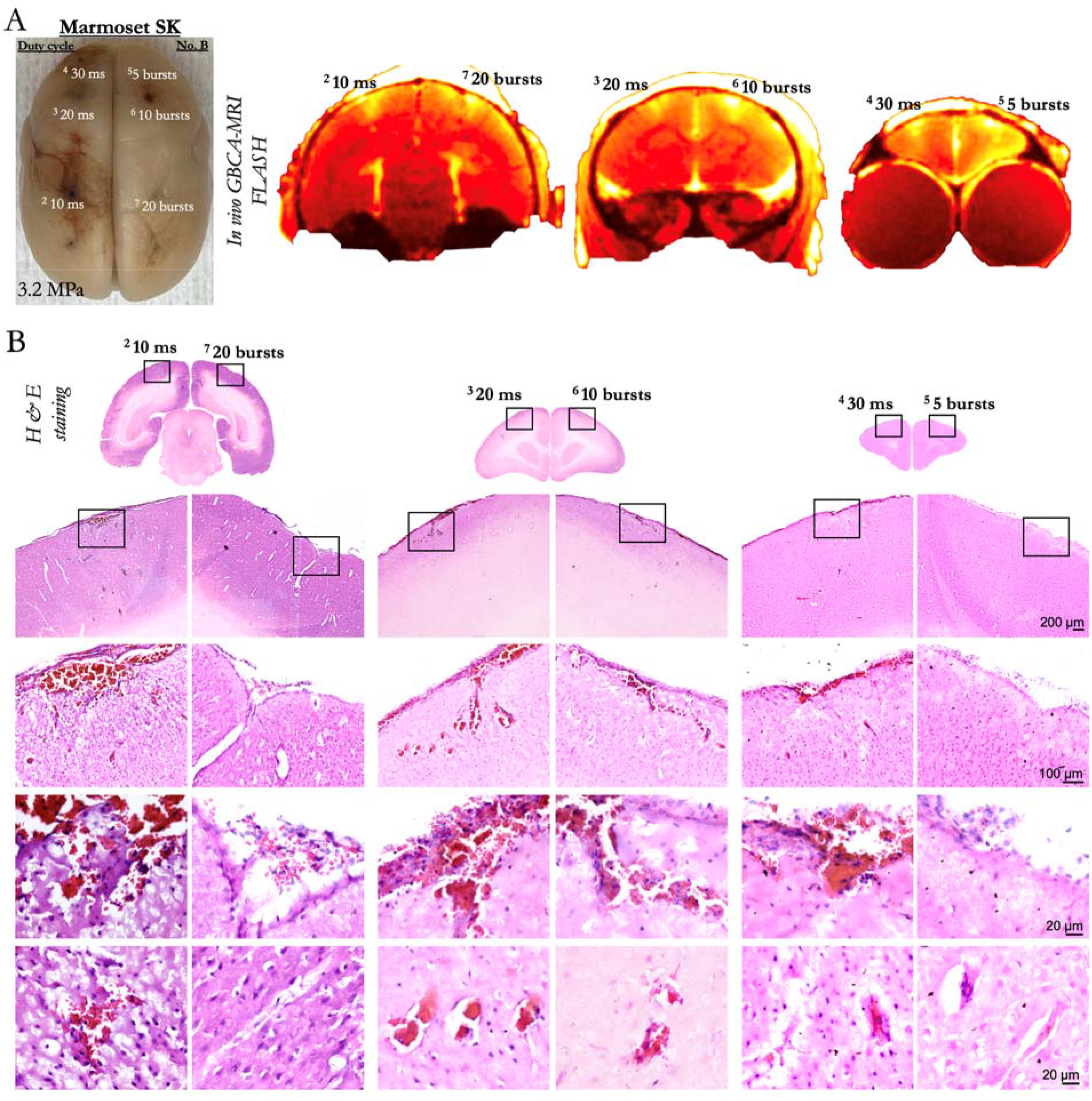
Cortical damage caused by high acoustic pressure and microbubble dose. (A) shows images derived from Figure 6 for reference for the sonication locations. (B) shows H & E staining at the sonicated locations, marked by the black boxes and zoomed, with scale bars at 200, 100, 20 μm.

## References

1. Pardridge, W. M. Drug Targeting to the Brain. Pharm. Res. 24, 1733–1744 (2007).

2. Hynynen, K., McDannold, N., Vykhodtseva, N. & Jolesz, F. A. Non invasive MR Imaging–guided Focal Opening of the Blood-Brain Barrier in Rabbits. Radiology 220, 640–646 (2001).

3. McDannold, N., Vykhodtseva, N. & Hynynen, K. Targeted disruption of the blood-brain barrier with focused ultrasound: Association with cavitation activity. Phys. Med. Biol. 51, 793–807 (2006).

4. McDannold, N., Arvanitis, C. D., Vykhodtseva, N. & Livingstone, M. S. Temporary disruption of the blood-brain barrier by use of ultrasound and microbubbles: Safety and efficacy evaluation in rhesus macaques. Cancer Res. 72, 3652–3663 (2012).

5. Pacia, C. P. et al. Feasibility and safety of focused ultrasound-enabled liquid biopsy in the brain of a porcine model. Sci. Rep. 10, 1–9 (2020).

6. Choi, J. J., Pernot, M., Small, S. A. & Konofagou, E. E. Noninvasive, transcranial and localized opening of the blood-brain barrier using focused ultrasound in mice. Ultrasound Med. Biol. 33, 95–104 (2007).

7. Karakatsani, M. E. M., Samiotaki, G. M., Downs, M. E., Ferrera, V. P. & Konofagou, E. E. Targeting Effects on the Volume of the Focused Ultrasound-Induced Blood-Brain Barrier Opening in Nonhuman Primates in Vivo. IEEE Trans. Ultrason. Ferroelectr. Freq. Control 64, 798–810 (2017).

8. Lapin, N. A., Gill, K., Shah, B. R. & Chopra, R. Consistent opening of the blood brain barrier using focused ultrasound with constant intravenous infusion of microbubble agent. Sci. Rep. 10, 1–11 (2020).

9. Kobus, T., Vykhodtseva, N., Pilatou, M., Zhang, Y. & McDannold, N. Safety Validation of Repeated Blood-Brain Barrier Disruption Using Focused Ultrasound. Ultrasound Med. Biol. 42, 481–492 (2016).

10. Ozdas, M. S. et al. Non-invasive molecularly-specific millimeter-resolution manipulation of brain circuits by ultrasound-mediated aggregation and uncaging of drug carriers. Nat. Commun. 11, 1–15 (2020).

11. Trinh, D. et al. Microbubble drug conjugate and focused ultrasound blood brain barrier delivery of AAV-2 SIRT-3. Drug Deliv. 29, 1176–1183 (2022).

12. Noroozian, Z. et al. MRI-Guided Focused Ultrasound for Targeted Delivery of rAAV to the Brain. Methods Mol. Biol. 1950, 177–197 (2019).

13. Meng, Y., Hynynen, K. & Lipsman, N. Applications of focused ultrasound in the brain: from thermoablation to drug delivery. Nat. Rev. Neurol. 17, 7–22 (2021).

14. Schaeffer, D. J. et al. Divergence of rodent and primate medial frontal cortex functional connectivity. Proc. Natl. Acad. Sci. U. S. A. 117, 21681–21689 (2020).

15. Sheikov, N., McDannold, N., Vykhodtseva, N., Jolesz, F. & Hynynen, K. Cellular mechanisms of the blood-brain barrier opening induced by ultrasound in presence of microbubbles. Ultrasound Med. Biol. 30, 979–989 (2004).

16. Chowdhury, S. M., Abou-Elkacem, L., Lee, T., Dahl, J. & Lutz, A. M. Ultrasound and microbubble mediated therapeutic delivery: Underlying mechanisms and future outlook. J. Control. Release 326, 75–90 (2020).

17. Ohl, C. D. et al. Sonoporation from jetting cavitation bubbles. Biophys. J. 91, 4285–4295 (2006).

18. Abrahao, A. et al. First-in-human trial of blood-brain barrier opening in amyotrophic lateral sclerosis using MR-guided focused ultrasound. Nat. Commun. 10, 4373 (2019).

19. Wang, S. et al. Non-invasive, Focused Ultrasound-Facilitated Gene Delivery for Optogenetics. Sci. Rep. 7, 1–7 (2017).

20. Ogawa, K. et al. Focused ultrasound/microbubbles-assisted BBB opening enhances LNP-mediated mRNA delivery to brain. J. Control. Release 348, 34–41 (2022).

21. Liu, C. et al. Marmoset Brain Mapping V3: Population multi-modal standard volumetric and surface-based templates. Neuroimage 226, 117620 (2021).

22. Schaeffer, D. J. et al. An open access resource for functional brain connectivity from fully awake marmosets. Neuroimage 252, 119030 (2022).

23. Paxinos, G., Watson, C., Petrides, M., Rosa, M. & Tokuno, H. The marmoset brain in stereotaxic coordinates. (‘Elsevier Academic Press, San Diego, 2012).

24. Hikishima, K. et al. Population-averaged standard template brain atlas for the common marmoset (Callithrix jacchus). Neuroimage 54, 2741–2749 (2011).

25. Liu, C. et al. A resource for the detailed 3D mapping of white matter pathways in the marmoset brain. Nat. Neurosci. 23, 271–280 (2020).

26. Paxinos, G., Watson, C., Petrides, M., Rosa, M. & Tokuno, H. The marmoset brain in stereotaxic coordinates. (Academic Press, 2012).

27. Bock, N. A., Paiva, F. F. & Silva, A. C. Fractionated manganese-enhanced MRI. NMR Biomed. 21, 473–478 (2008).

28. Schindelin, J. et al. Fiji: An open-source platform for biological-image analysis. Nat. Methods 9, 676–682 (2012).

29. Yeh, F. C., Verstynen, T. D., Wang, Y., Fernández-Miranda, J. C. & Tseng, W. Y. I. Deterministic diffusion fiber tracking improved by quantitative anisotropy. PLoS One 8, e80713 (2013).

30. O’Reilly, M. A., Muller, A. & Hynynen, K. Ultrasound Insertion Loss of Rat Parietal Bone Appears to Be Proportional to Animal Mass at Submegahertz Frequencies. Ultrasound Med. Biol. 37, 1930–1937 (2011).

31. Ferrara, K., Pollard, R. & Borden, M. Ultrasound microbubble contrast agents: Fundamentals and application to gene and drug delivery. Annu. Rev. Biomed. Eng. 9, 415–447 (2007).

32. Daneman, R. & Prat, A. The blood-brain barrier. Cold Spring Harb. Perspect. Biol. 7, a020412 (2015).

33. Kovacs, Z. I. et al. Disrupting the blood-brain barrier by focused ultrasound induces sterile inflammation. Proc. Natl. Acad. Sci. U. S. A. 114, E75–E84 (2017).

34. Jordão, J. F. et al. Amyloid-β plaque reduction, endogenous antibody delivery and glial activation by brain-targeted, transcranial focused ultrasound. Exp. Neurol. 248, 16–29 (2013).

